# Bioengineering an *in situ* ovary (ISO) for fertility preservation

**DOI:** 10.1101/2020.01.03.893941

**Authors:** Michael J. Buckenmeyer, Meena Sukhwani, Aimon Iftikhar, Alexis L. Nolfi, Ziyu Xian, Srujan Dadi, Zachary W. Case, Sarah R. Steimer, Antonio D’Amore, Kyle E. Orwig, Bryan N. Brown

**Affiliations:** Department of Bioengineering, University of Pittsburgh, Pittsburgh, PA, USA; McGowan Institute for Regenerative Medicine, University of Pittsburgh, Pittsburgh, PA, USA; Department of Obstetrics, Gynecology and Reproductive Sciences, Magee-Womens Research Institute, University of Pittsburgh School of Medicine, Pittsburgh, PA, USA; Fondazione RiMED, Via Bandiera 11, 90133 Palermo, Italy

## Abstract

Female cancer patients who have undergone chemotherapy have an elevated risk of developing premature ovarian failure (POF) and infertility. Experimental approaches to treat iatrogenic infertility are evolving rapidly; however, there remain challenges and risks that have hindered clinical translation. To address these concerns, we developed an ovarian-specific extracellular matrix hydrogel to facilitate follicle delivery and establish an *in situ* ovary (ISO). We demonstrate sustainable engraftment, natural pregnancy and the birth of healthy pups after intraovarian microinjection of isolated exogenous follicles in a chemotherapy-induced POF mouse model. Our results suggest that the methods described could offer a minimally invasive alternative for the restoration of the ovarian reserve post-chemotherapy.

---

Ovarian follicles are the major functional component of the ovary that produce hormones (e.g., estrogen) and mature eggs for ovulation^1,2^. Chemotherapy or radiation treatments for cancer or other conditions can deplete the ovarian follicle pool, which can result in premature ovarian failure (POF), compromising ovarian hormone production and fecundity^3–5^. Ovarian tissue cryopreservation is an experimental option used to preserve the fertility of patients who cannot afford to delay gonadotoxic treatment. Upon remission, the ovarian cortex, which is rich in primordial follicles, can be transplanted back into patient survivors and ovulate eggs to establish natural pregnancies or be stimulated to produce eggs for *in vitro* fertilization (IVF)^6–14^. However, ovarian tissue transplantation is an invasive surgical procedure and may not be appropriate in cases where there are concerns that ovarian tissues may harbor malignant cells (e.g., leukemia).

Artificial ovaries, coupling follicle isolation with implantable biomimetic scaffolds, have been proposed as a method to reduce the risk of malignant cell transplantation while also aiming to restore fertility. However, successful applications of this approach have solely used pre-formed constructs^15–18^, which would still require an invasive implantation procedure. Here we report a strategy to reduce the surgical burden by establishing an *in situ* ovary (ISO) through the intraovarian microinjection of isolated immature follicles facilitated by a porcine ovarian extracellular matrix (OECM) hydrogel. Follicle transplant recipient chemotherapy-induced POF (ciPOF) mice were bred to fertile males and produced donor follicle (GFP+)-derived progeny.

## Results

### Biomaterial selection and tissue processing

The damaging effects of chemotherapy on ovarian tissues significantly reduces a patient’s follicle population, which has a direct impact on fertility and endocrine function^3–5^. In addition, chemical treatments have been implicated in microvasculature and stromal cell irregularities culminating in a compromised environment for cell survival^3^. These unfavorable conditions cause a depletion of ovarian follicles and may reduce follicle viability post-transplantation^16,19^. Therefore, to re-establish the ovarian tissue microenvironment and repopulate the depleted endogenous follicle pool, we have bioengineered an ISO using an OECM hydrogel to facilitate intraovarian follicle transplant and provide a temporary niche to aid follicle engraftment and survival (Fig. 1a).

**Fig. 1.**
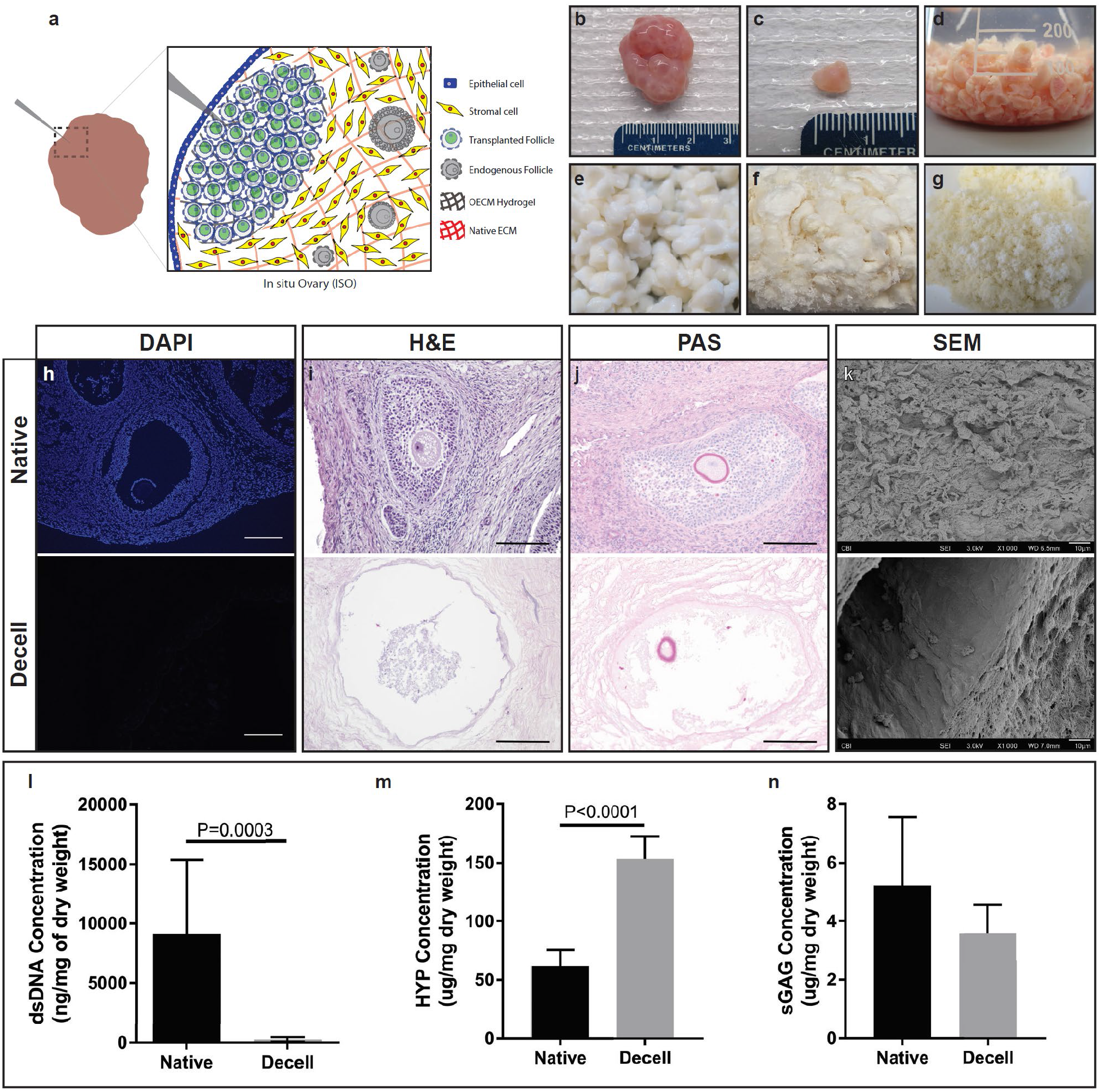
Central hypothesis, decellularization process and tissue characterization.

**a**, The graphical representation shows our hypothesis for restoring fertility in patients with chemotherapy-induced premature ovarian failure (ciPOF). Intraovarian microinjection of an ovarian-specific ECM (OECM) hydrogel can support the delivery and long-term survival of exogenous primordial follicles (green) within an *in situ* ovary (ISO). The damaged ovarian tissue primarily consists of stromal cells (yellow) and a depleted population of endogenous follicles (gray). **b**, Young (< 1 year old) porcine ovaries were sourced for decellularization. **c**, Ovaries were diced into small cubes (~0.125 cm^3^). **d**, Cubed ovaries were added to a flask and decellularized using enzymes and detergents. **e**, Decellularized ovarian tissues appeared white. **f**, Ovaries were frozen then lyophilized to remove their water content. **g**, Powdered OECM was prepared using a mill. **h-j**, Native (top row) and decellularized (bottom row) images of DAPI (200 μm scale), H&E (100 μm scale), and periodic acid-Schiff (PAS) (100 μm scale) staining determined that decellularized tissues removed cellular content while preserving ovarian tissue morphology. **k**, Scanning electron micrographs (SEM) show a dense cellular ultrastructure in native ovaries in comparison to a porous decellularized scaffold (10 μm scale). **l**, PicoGreen assay indicated that decellularized ovarian tissues significantly reduced the dsDNA concentration. Data represent mean ± s.e.m. of ng/mg dry weight. P values by unpaired, two-tailed t-test. **m**, Hydroxyproline (HYP) concentration was significantly enriched in decellularized tissues and **n**, sulfated-Glycosaminoglycans (sGAG) levels did not differ significantly between native and decellularized samples. Data represent mean ± s.e.m. of μg/mg dry weight. P values by unpaired, two-tailed t-test.

To prepare acellular ovarian scaffolds, porcine ovaries (Fig. 1b) were diced (Fig. 1c) then processed using a series of enzymatic and detergent washes to remove immunogenic material (Fig. 1d). Trypsin and EDTA were used in tandem to disrupt cell adhesions to the ECM prior to treatments with Triton X-100 and sodium deoxycholate, which act to permeabilize cell membranes. Tissues transitioned from an initial opaque appearance to translucent at the conclusion of the decellularization steps (Fig. 1e). Intermediate water washing steps proved to be critical for the complete removal of residual cells and detergent from the tissues. Once the tissues were completely decellularized they were frozen, lyophilized (Fig. 1f) and milled into a powder (Fig. 1g) prior to biochemical testing and downstream processing.

To demonstrate the effective removal of immunogenic components and preservation of extracellular matrix (ECM) components, we performed a set of histological stains and biochemical assays. Fluorescence staining with 4’,6-diamidino-2-phenylindole (DAPI) showed few, if any nuclei present within the decellularized tissues in comparison to native ovarian tissue controls (Fig. 1h). H&E (Fig. 1i) and PAS (Fig. 1j) staining showed a clear retention of ovarian microarchitecture, such as structural aspects of follicles, zona pellucida and corpora lutea, while sparse cellular content was visible. This highlights the effectiveness of our method to successfully remove cellular content, while causing limited disruption of tissue-specific morphology. Scanning electron micrographs (SEM) further detailed the dense cellular content within native ovarian tissues, whereas decellularized tissues appeared to show vacated follicular compartments surrounded by a porous scaffold (Fig. 1k). A PicoGreen assay demonstrated greater than 98% reduction of dsDNA between native (9126 ± 1988 ng/mg) and decellularized (262.4 ± 59.96 ng/mg) samples (Fig. 1l). Gel electrophoresis further showed a lack of DNA (Supplementary Fig. 1) within the decellularized tissues in comparison to native controls, suggesting a reduced potential for disease transmission and adverse immune reaction to cellular contents. Collagen and sulfated glycosaminoglycans (sGAG) were also examined to determine their retention post-decellularization. A hydroxyproline (HYP) assay was used to estimate the total collagen content within the scaffold. Native tissues (61.95 ± 6.064 ug/mg) contained significantly less HYP as a percentage of dry weight than decellularized tissues (153.3 ± 8.564 ug/mg) due to the loss of cellular mass; however, under this assumption the total collagen content within the decellularized scaffold as a fraction of the dry weight of all components was enriched after decellularization (Fig. 1m). sGAG content was also preserved with no significant difference observed between native (5.24 ± 1.03 ug/mg) and decellularized (3.59 ± 0.436 ug/mg) samples (Fig. 1n).

### Ovarian tissue specificity present post-decellularization

The ECM is composed of a tissue-specific milieu of secreted proteins and proteoglycans that support the desired functions of a tissue^20–22^. In the ovary, the ECM undergoes dynamic remodeling throughout the reproductive life span and is essential for regulating folliculogenesis and ovulation^23,24^. Specifically, OECM provides mechanical support, maintains normal cell morphology, promotes cell proliferation and steroidogenesis^24^. Additionally, the OECM can sequester hormones and growth factors within the follicle niche to facilitate paracrine and endocrine signaling^24,25^. Therefore, the retention of OECM proteins would be ideal for supporting follicles within the ISO. To determine the effects of decellularization on ECM retention, we characterized a subset of the most highly expressed OECM proteins: Collagen I, Collagen IV, laminin and fibronectin^23,24,26,27^.

Immunohistochemistry revealed that Collagen I was distributed uniformly in the native samples, with a slight enrichment surrounding the thecal compartments of the follicles in decellularized samples (Fig. 2a). Collagen IV was also labeled, showing definitive staining within the basement membrane of the epithelial layer and the basal lamina of individual follicles in both the native and decellularized groups (Fig. 2b). Similar to Collagen IV, laminin was predominantly found within the basal lamina adjacent to the theca interna surrounding follicles (Fig. 2c). Finally, fibronectin appeared to be conserved throughout the ovarian tissues with little to no differences in distribution noted between the native and decellularized groups (Fig. 2d).

**Fig. 2.**
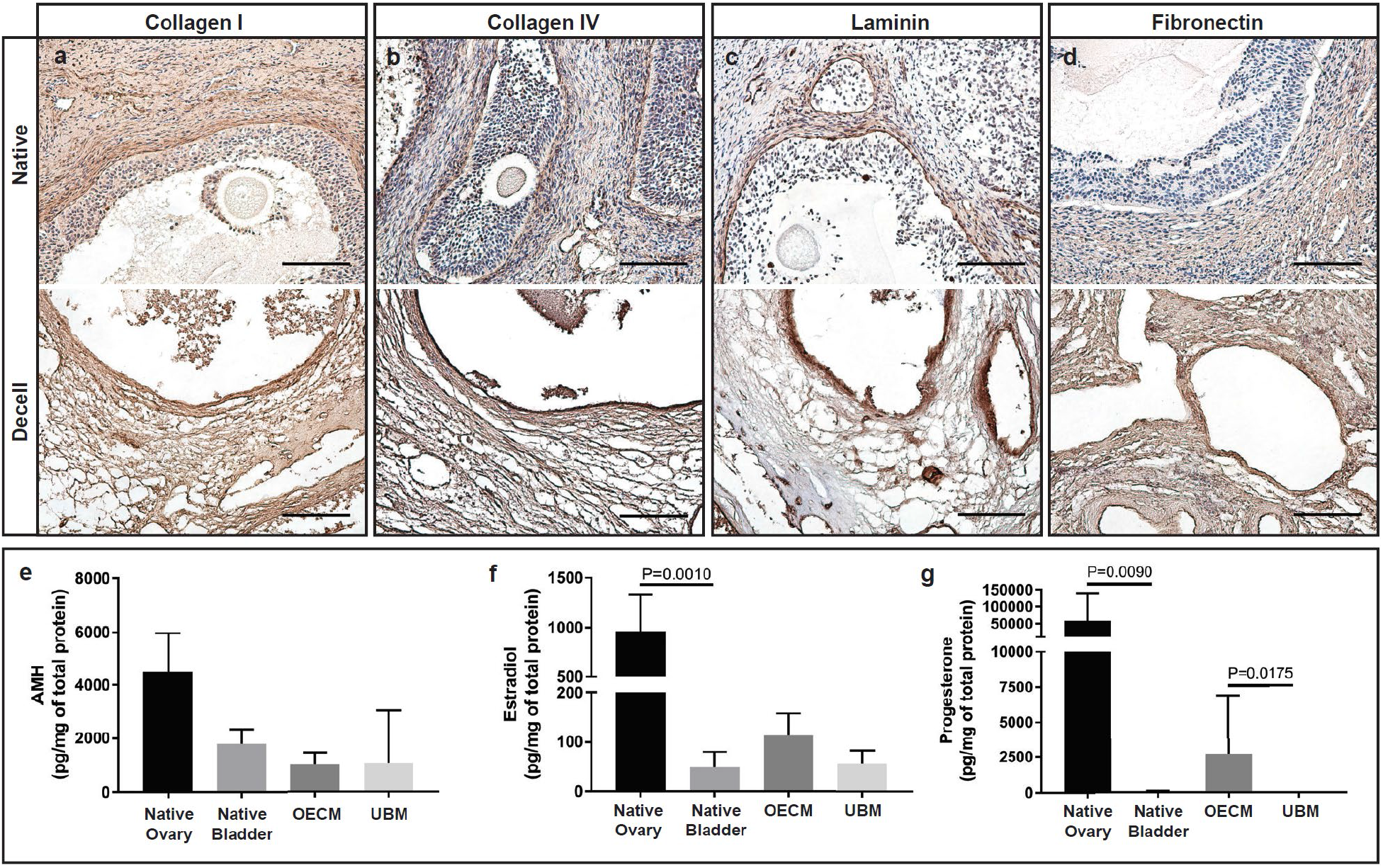
Preservation of ovarian-specific ECM proteins and reproductive hormones.

Immunohistochemistry (IHC) images of native (top row) and decellularized (bottom row) ovarian tissues shows the distribution of extracellular matrix proteins: **a**, Collagen I was uniformly expressed throughout each of the tissues. **b**, Collagen IV appeared to concentrate in the basal lamina of the follicles and the basement membrane surround the epithelial layer of the ovary. **c**, Laminin was present within the thecal compartment and **d**, Fibronectin was evenly expressed in lower concentrations; however, antibody staining was enriched in decellularized tissues. Scale, 100 μm. Ovarian hormones secreted by follicular cells were quantified from total protein using enzyme-linked immunosorbent assays (ELISAs). **e**, Anti-Müllerian hormone (AMH) was measured at high concentrations within native ovarian tissues but was reduced by >50% within decellularized OECM. **f**, Estradiol concentrations were significantly higher in native ovary in comparison to native bladder and OECM was two-fold greater than UBM. **g**, Progesterone levels were significantly greater in both native ovary and OECM in comparison to native bladder and UBM, respectively. ELISAs were conducted using 100 ovaries batched into 5 independent samples (n = 20 ovaries). Data represent mean ± s.e.m. of pg/mg of total protein. P values by one-way ANOVA using Kruskal-Wallis with Dunn’s multiple comparisons test. Native (ovary and bladder) and decellularized groups (OECM and UBM) were compared separately.

Ovarian hormones and growth factors sequestered in the OECM orchestrate both local and systemic endocrine function. The hypothalamic-pituitary-gonadal (HPG) axis stimulates the production of ovarian hormones, which act to modulate hormone production in a cyclic manner^1^. The hypothalamus produces gonadotropin releasing hormone (GnRH), which stimulates the secretion of follicle stimulating hormone (FSH) and luteinizing hormone (LH)^1^. FSH and LH trigger the production of estradiol, follicle development and ovulation. Estradiol from the ovulatory follicle and progesterone from the resulting corpus luteum provide feedback to either inhibit or stimulate hormone secretion from the hypothalamus and pituitary^1^. Spatiotemporal production of these reproductive hormones primarily facilitates follicle development, ovulation and pregnancy^1^.

For this study, we were particularly interested in hormones produced by the ovary due to their roles in follicle selection. Specifically, anti-Müllerian hormone (AMH), estradiol and progesterone. AMH is produced by granulosa cells of pre-antral and antral follicles^28^. As AMH levels increase, it can inhibit the recruitment of primordial follicles and decrease the responsiveness of large pre-antral/antral follicles to FSH. AMH is one of the few hormones that are produced during the early stages of folliculogenesis, which are widely recognized as gonadotropin-independent^29^. Estradiol is also produced by follicular cells and is most commonly known for its role in the LH surge which triggers ovulation; however, at low concentrations, estradiol can function as a negative regulator of FSH, which inhibits follicle growth^29,30^. The corpus luteum, which arises from the cells of ovulatory follicles and is present during the late luteal phase, produces high levels of progesterone, which is necessary for maintaining pregnancy. Like estradiol, progesterone can also inhibit FSH production further delaying follicle growth^29^.

We hypothesized that these ovarian hormones may be important for the survival of transplanted immature follicles within the ISO. Therefore, to elucidate the effects of decellularization on the disruption of these components, we processed a large batch of 100 ovaries separated into five groups (n = 20 per group) analyzed as independent samples. As controls, we used both native and decellularized urinary bladder matrix (UBM), collected from female pigs and prepared as previously described^31^, to determine if there were significant differences in the hormone concentrations based upon tissue source. After tissue homogenization, protein was extracted from each group and tested using biochemical assays for AMH, estradiol and progesterone. ELISA quantification determined that decellularized OECM samples contained low concentrations of each of the ovarian-specific analytes: AMH (1031 ± 192.9 pg/mg), estradiol (113.8 ± 19.63 pg/mg) and progesterone (2697 ± 1890 pg/mg) (Fig. 2e-g). Native ovaries contained significantly higher levels of estradiol and progesterone when compared to native bladder. Furthermore, decellularized OECM had significantly higher progesterone values than UBM. Additional analytes associated with follicle development, insulin-growth factor (IGF-1) and vascular endothelial growth factor (VEGF), were also tested but were undetectable within the decellularized samples (data not shown).

### Ovarian hydrogel properties modified with changes in ECM concentration

Once the decellularized tissues were processed and characterized, we solubilized the OECM then neutralized the digested material to prepare the hydrogels. Visibly transparent hydrogels were formed after exposure to physiological conditions for approximately 20 minutes (Fig. 3a). We used SEM to evaluate the hydrogel ultrastructural properties at 4 and 8 mg/mL ECM concentrations (Fig. 3b,c). Fiber network characteristics were quantified using SEM imaging and digital image analysis^32^ with concentration dependent effects observed with a significant increase in fiber diameter, fiber length and bulk porosity in the 8 mg/mL OECM hydrogel group (Fig. 3d-h). These results indicate that individual fiber and large-scale network properties such as the bulk porosity are dependent on ECM concentration.

**Fig. 3.**
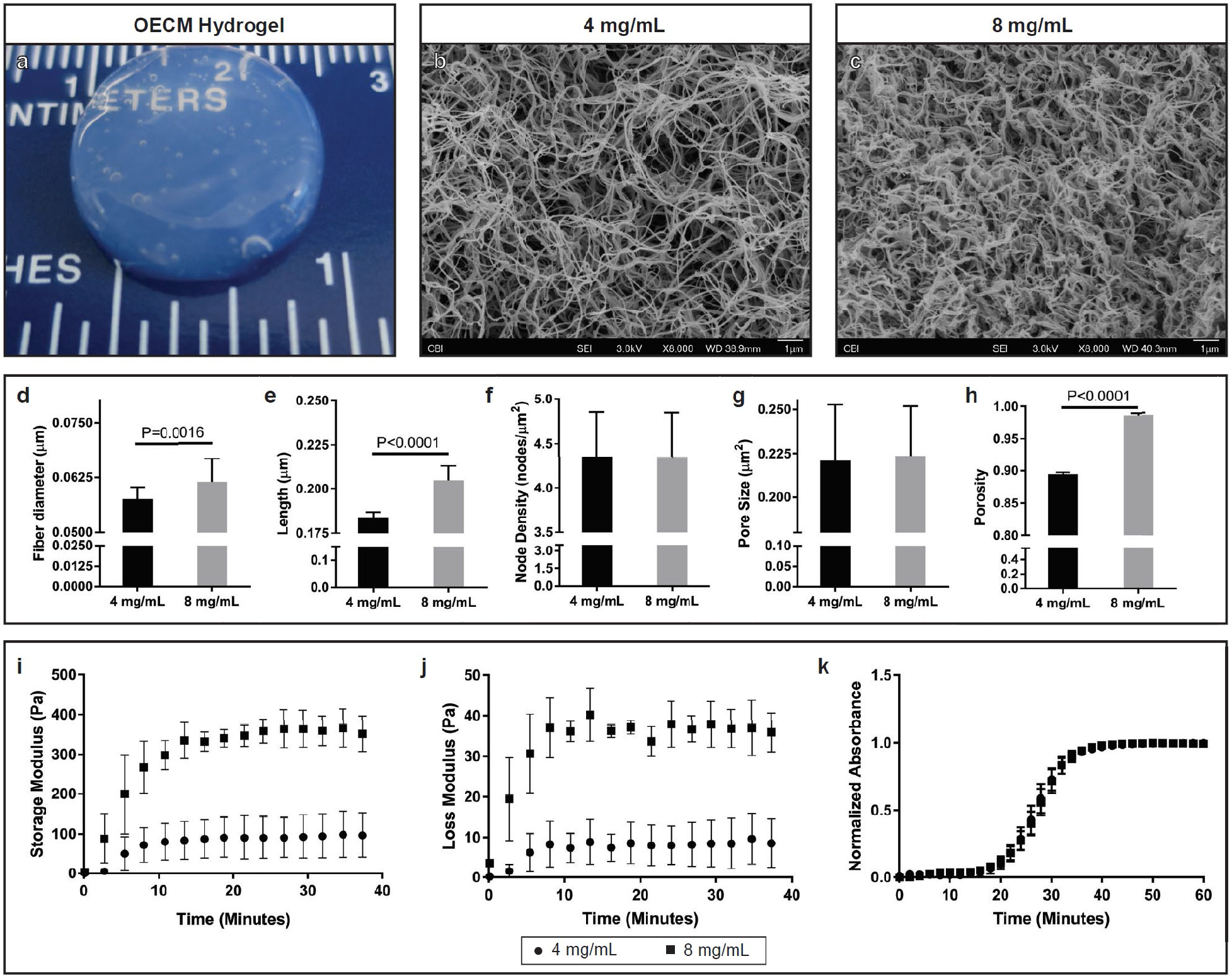
OECM hydrogel preparation, characterization of ultrastructure and viscoelastic properties.

**a**, Solubilized ovarian ECM (OECM) formed transparent hydrogels upon neutralization and exposure to physiologic conditions. **b-c**, Scanning electron micrographs (SEM) show a fibrous and porous architecture at both 4 and 8 mg/mL OECM concentrations. **d-h**, Hydrogel fiber network characteristics were quantified using a Matlab algorithm with significant increases in fiber diameter, fiber length and porosity directly correlating with an increase in OECM concentration. Data represent mean ± s.e.m. for all parameters. P values by unpaired, two-tailed t-test. Viscoelastic material properties were quantified using a rheological time sweep. **i**, Storage (G’) and **j**, Loss (G”) moduli increased dramatically with the higher 8 mg/mL OECM concentration. **k**, Turbidimetric gelation kinetics showed that gelation was conserved with a change in OECM concentration.

To determine the viscoelastic properties of the OECM hydrogel, we performed a rheological time sweep on varying ECM concentrations. An increase in ECM concentration from 4 to 8 mg/mL appeared to correlate with an elevated storage (G’) and loss (G”) moduli (Fig. 3i,j). However, there were no observable differences in turbidimetric gelation kinetics with a change in ECM concentration (Fig. 3k). Gelation time varied based upon the test conditions. Specifically, direct conduction with the Peltier plate achieved complete gelation approximately 15 minutes prior to the samples heated via convection during gelation kinetics testing. However, once gelation initiated, hydrogels from both test formats consistently solidified within 20 minutes.

### Alkylating agents significantly reduce endogenous follicle population

After developing and characterizing the OECM hydrogel as a carrier for follicle injection, we created a clinically relevant ciPOF mouse model. We chose to induce POF using alkylating agents, busulfan and cyclophosphamide, due to their known cytotoxic effects on ovarian tissues and cells^3,5,33,34^. Briefly, a single intraperitoneal (IP) injection was given to 6-week old nude female mice. Dosing was titrated to determine an appropriate treatment that would significantly reduce the endogenous follicle pool to lower or eliminate the chances of fertility. The following doses were tested, abbreviated as busulfan-cyclophosphamide (mg/kg): (1) 12-100 (2) 12-200 (3) 24-100 (4) 24-200.

At three weeks post-IP injection, histological staining with Weigert’s Hematoxylin-Picric Methyl Blue clearly illustrated the damaging effects of each chemotherapy regimen on the follicle population within the ovaries. Dose dependent effects were observed with elevated levels of busulfan and cyclophosphamide reducing both follicle quantity and apparent tissue volume (Fig. 4a-e). Follicle quantification supported these observations as increasing dosage confirmed a decreasing total follicle number (Fig. 4f). Follicles were counted based upon developmental stage (Supplementary Fig. 2). Primordial follicles were especially sensitive to these treatments with a pronounced reduction in this immature stage in comparison to more developed follicles (Fig. 4g). After comparing the effects of each treatment, we determined that the 24-100 group offered both a significant reduction of the primordial follicle population (approximately 94%) without appreciably compromising the ovarian tissue volume. Therefore, this group was selected as the primary treatment to prepare the ciPOF follicle recipient and control mice.

**Fig. 4.**
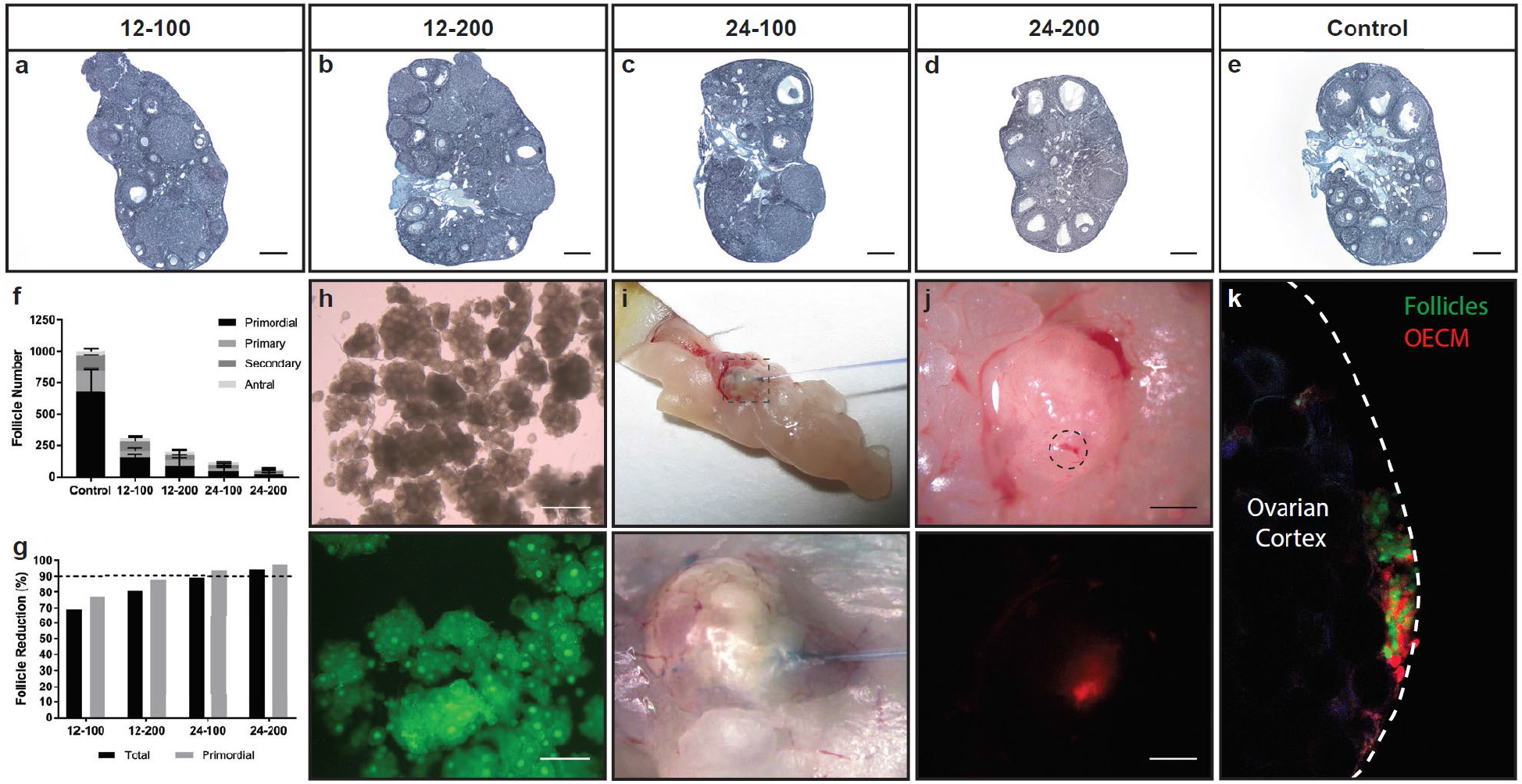
surface of the ovarian epitheliumSelection of chemotherapy dose and confirmation of an established *in situ* ovary (ISO).

Four doses of busulfan-cyclophosphamide **a**, 12-100 mg/kg **b**, 12-200 mg/kg **c**, 24-100 mg/kg **d**, 24-200 mg/kg **e**, non-injected (control) were administered via single IP injection. Ovaries were excised at 3 weeks post treatment and stained using Weigert’s Hematoxylin Picric Acid Methyl Blue. Scale, 250 μm. **f**, Follicles were manually counted, quantified and classified by developmental stage showing a steady decline of the total follicle number with increasing dose. Additionally, busulfan appeared to have an enhanced effect on follicle depletion in comparison to cyclophosphamide. **g**, Primordial follicle reduction was >90% (dotted line) in both the 24-100 and 24-200 indicative of a severe depletion of the ovarian reserve, significantly lowering potential fertility and could represent premature ovarian failure (POF). **h**, Intact GFP follicles were enzymatically isolated using Liberase TM and imaged under bright field (top) and fluorescence (bottom). Scale, 500 μm. **i**, Brightfield images show the gross morphology of the ovary during microinjection (top) and magnified to show the injection site for follicle transplant (bottom). **j**, Pressurized microinjection was tested as a potential technique to deliver the OECM hydrogel (TRITC-labeled) and visualized under bright field (top) and fluorescence (bottom). Scale, 500 μm. **k**, Multiphoton images confirmed that intraovarian follicle microinjection of the isolated GFP follicles (green) and OECM (red) co-localized within the ovarian cortex to form an *in situ* ovary. The dotted line (white) indicates the outer surface of the ovarian epithelium.

### Enzymatic follicle isolation and microinjection provides an efficient transplant procedure

Follicle incubation time from isolation to transplant is a concern for cell therapy applications as it can directly impact viability^35^. Therefore, we adapted an existing enzymatic isolation protocol^36^ to reduce the time needed to obtain a large pool of immature follicles for transplant. To enable identification of transplanted versus endogenous follicles, we isolated follicles from transgenic mice exhibiting ubiquitous GFP under the chicken β-actin (CAG) promoter. We tested enzyme concentrations of 10% and 20% for both Liberase TM and DH (13 Wünsch units/mL) to determine their effects on follicle disaggregation and quality. Briefly, Liberase TM or DH was added to the minced ovaries then two five-minute cycles of physical agitation at 37°C were performed with a minute of pipetting after each cycle. After assessing each sample, we determined that the best formulation was the 10% Liberase TM, which released a large population of morphologically normal GFP follicles during the 12-minute isolation procedure (Fig. 4h). In order to estimate the number of follicles isolated with this procedure, follicles were manually counted using a hemocytometer. There were approximately 1.5×10^3^ total follicles isolated per ovary with 74.4% of this population identified at the primordial stage.

Once we showed that it was feasible to efficiently obtain follicles, we wanted to test the efficacy of using microinjection to transplant follicles into the ovarian cortex of ciPOF mice. Immature follicles naturally reside within the ovarian cortex, as this region of the ovary has mechanically distinct properties that support the early stages of folliculogenesis^19,37^. Therefore, it was important to identify a cell delivery modality that would precisely dispense follicles into or near the cortex. Previously, intragonadal cell delivery has been shown successfully using microinjection^38^. So, we decided to implement this technique to facilitate intraovarian follicle transplant to establish an ISO. First, we tested the delivery of the OECM hydrogel alone via microinjection. A small volume of hydrogel was injected into the ovarian cortex, and the recipient animal was sacrificed to visualize the injection site (Fig. 4i). Bright field images clearly illustrated Trypan blue dye at the site of injection at the tissue surface (Fig. 4j). Furthermore, the use of a TRITC-labeled OECM hydrogel allowed us to identify the hydrogel at the ovarian surface post-injection (Fig. 4j). These results confirmed this method as a suitable delivery mechanism for the viscous OECM within a specific anatomical location of the ovary. Finally, we performed the same experiment with the addition of isolated GFP follicles to determine if the gel and follicles could be delivered simultaneously, resulting in the formation of an ISO. Ovaries excised from the TRITC-OECM hydrogel and GFP follicle microinjection clearly indicated a co-localization of the OECM and follicles within the ovarian cortex (Fig. 4k).

### Microinjected follicles give rise to multiple generations of GFP pups

We used our ciPOF model to investigate the therapeutic potential of an ISO to support follicle survival after intraovarian microinjection. Approximately 1.0×10^3^ GFP+ follicles resuspended in 8 μL of OECM hydrogel were microinjected into the ovarian cortex of ciPOF nude female mice (n = 2). Non-injected ciPOF nude female mice were used as controls (n = 2). To reduce tissue damage due to needle puncture, only a single follicle injection was performed on each ovary. Freshly isolated GFP follicles from 6-14 day female (DBA-GFP/nu-) mice were used to ensure a predominantly immature follicle population at the time of injection.

To determine the effects of follicle microinjection on fertility, follicle recipient and non-injected control ciPOF nude female mice were mated with nude male mice for three breeding cycles (~100 days). The breeding strategy was designed to definitively distinguish between pups derived from transplanted or endogenous follicles (Supplementary Fig. 3a). Pups from endogenous follicles would be nude, lacking fur with red eyes, whereas pups from transplanted GFP follicles would have fur with dark eyes and a 50% chance of glowing green (GFP/nu-). Breeding resulted in consecutive litters containing GFP+/nu-pups from one of the follicle recipient mice (Supplementary Fig. 3b). The first breeding cycle yielded one GFP pup (GFP/nu-001) out of three healthy offspring followed by a single GFP pup (GFP/nu-002) during the second cycle (Fig. 5a,b). The other follicle transplant recipient did not produce any offspring throughout the mating period. Only one of the control mice produced multiple litters with 11 pups during the first cycle and 5 pups during the second cycle with no GFP expression. None of the mice produced any litters during the third breeding trial. As a chemotherapy dosing control, ciPOF nude female mice treated with a less severe dosage (12-100 mg/kg) were mated and produced a single GFP pup (GFP/nu-003) during their third mating cycle (Fig. 5c), which provides additional evidence for long-term follicle viability post-transplant (Supplemental Fig. 3b). A follow-up mating study was performed to test the reproductive health of the GFP pups generated from follicle transplantation. The GFP offspring resulting from follicle transplant (DBA-GFP+/nu-) were bred with DBA wild-type (GFP-/nu+) (Supplementary Fig. 3c) and inbred with each mating pair (Supplementary Fig. 3d) producing multiple large litters of hemizygous (GFP^+/-^) and homozygous (GFP^+/+^) genetic backgrounds (Fig. 5d and Supplementary Fig. 3e). In each of the litters, GFP expression was clearly observed in the presence of a UV lamp and confirmed with genotyping (Fig. 5f).

**Fig. 5.**
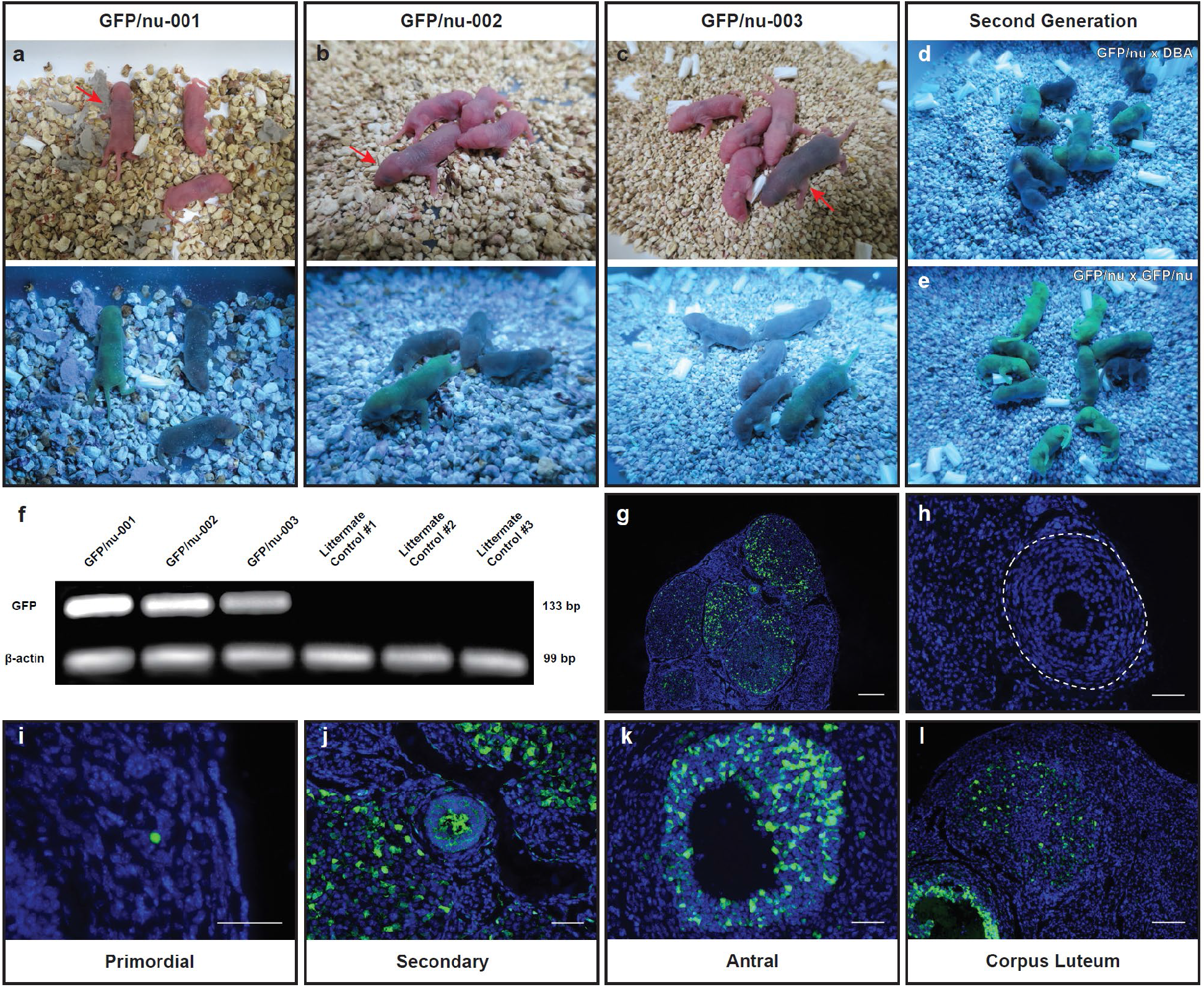
Intraovarian transplant gives rise to GFP offspring and supports follicle longevity.

The efficacy of intraovarian follicle transplant for fertility preservation was tested by injecting a pool of isolated GFP follicles within an OECM hydrogel into a ciPOF nude female mouse (24-100 mg/kg busulfan-cyclophosphamide). The follicle recipient mice were then mated to male mice of the same genetic background (nu/nu) to distinguish between progeny derived from exogenous (GFP^+^) or endogenous (GFP^-^) follicles. **a-b**, Two GFP pups (GFP/nu-001 and GFP/nu-002) were born in consecutive liters from the same mother as a direct result of the follicle transplantation. **c**, Another GFP pup (GFP/nu-003) was derived from a ciPOF female mouse (12-100 mg/kg busulfan-cyclophosphamide) during the third mating cycle (>100 days post-transplant). This demonstrates that an ISO can support injected follicles in a dose-independent manner and have long-term viability post-injection. Multiple litters of second generation pups were derived from both **d**, outbred (GFP/nu x DBA) and **e**, inbred (GFP/nu x GFP/nu) mice, which indicates that intraovarian follicle transplantation did not disrupt reproductive development. **f**, Genotyping of the GFP pups and littermate controls was confirmed using standard PCR and gel electrophoresis. GFP bands were only appeared within the GFP mouse samples and none within the littermate samples. β-actin was used as an internal control appearing in each of the samples tested. Gels were cropped and processed to highlight the bands of interest. Ovarian tissues were excised from ciPOF follicle recipient mice after three breeding cycles (106 days on average). Immunofluorescence staining was performed using DAPI (endogenous cells) and GFP (transplanted cells) to evaluate follicle survival. **g**, A representative image of GFP^+^ follicles within the transplanted ovary suggests that the transplanted cells integrated within the tissues and were actively developing. Scale, 200 μm. **h**, Presence of an endogenous secondary follicle (GFP^-^), indicated by dotted line (white). Scale, 50 μm. Various stages of follicle development were also present among the transplanted tissues, including: **i**, Primordial (Scale, 50 μm) **j**, Secondary (Scale, 50 μm) **k**, Antral (Scale, 50 μm) and **l**, Corpus Luteum (Scale, 100 μm).

### Intraovarian microinjection supports follicle longevity post-transplant

Finally, we wanted to determine the effects of this therapy on follicle longevity post-transplant. To answer this question, we performed immunofluorescent labeling of ovaries excised from ciPOF nude female mice after three breeding cycles (~100 days). Comprehensive imaging demonstrated significant GFP expression throughout the transplanted tissues and suggests that multiple follicles remained viable post-transplantation (Fig. 5g). Transplanted ovaries also retained growing endogenous follicles, which may additionally be supported by the ISO components (Fig. 5h). Non-injected control tissues lacked GFP expression and endogenous follicles appeared to be reduced in comparison to ovaries from transplanted mice (Supplemental Fig. 4). However, we could not confirm any definitive differences between residual endogenous follicles based solely upon the immunofluorescence images. Finally, we were able to identify follicles at various stages of development including primordial (Fig. 5i), secondary (Fig. 5j), antral (Fig. 5k), and corpus luteum (Fig. 5l), which indicates that the ISO can facilitate long-term follicle integration, maintenance, and development.

## Discussion

Women who cryopreserve oocytes or embryos prior to gonadotoxic treatments can use assisted-reproductive technologies (ART), such as IVF and embryo transfer to start a family^39,40^. To preserve eggs or embryos, the patient must first undergo hormone stimulation to collect mature oocytes. Controlled ovarian stimulation requires two or more weeks and is not a viable option for patients who have not reached reproductive maturity or who cannot afford to postpone treatment^41–43^. For these women, cryopreservation of intact ovarian tissues prior to treatment is the only potential option to naturally restore endocrine function and fertility. The current gold standard for fertility preservation in patients in remission is the autologous surgical transplantation of cryopreserved ovarian cortical strips^6,8,11–14,44–47^. To date, there have been numerous successful procedures performed in humans, resulting in greater than 130 live-births; however, the efficiency of this method remains low, with live-birth rates ranging from 23-36% of patients achieving live-births^48,49^. Although ovarian tissue transplantation has shown promise, it is an invasive procedure and carries a potential risk of reintroducing malignant cells back into the body^50^. To address these concerns, several pre-clinical experimental approaches have been proposed, including *in vitro* follicle maturation (IVM), the development of an artificial ovary and stem cell transplantation (Table 1).

**Table 1.** Summary of notable pre-clinical experimental therapies and their future outlook for fertility preservation. Several therapeutic approaches have been explored to ameliorate the effects of infertility. In general, these can be divided into three main categories: (1) Ovarian engraftment (2) Biomaterial-facilitated and (3) Intraovarian transplantation methods. Ovarian engraftment can be equated to cortical strip autotransplantation, which is currently considered as the gold-standard in the field. Biomaterial-facilitated methods have recently progressed toward the development of a functional artificial ovary, which uses a scaffold or hydrogel to support follicle transplantation. Intraovarian transplantation has been primarily focused on the delivery of female germline stem cells. Each of the in vivo studies listed have used varied models of infertility, induced either by ovariectomy or chemotherapy, with a limited number showing the ability to produce pups (highlighted in green). However, the clinical translation of these approaches may be hindered by invasive surgical procedures and *ex vivo* cell manipulation. To overcome these barriers to the clinic, we have proposed a minimally invasive technique to facilitate the delivery and support of ovarian follicles within an *in situ* ovary (ISO), which promotes a natural solution for restoring fertility.

| Approach | Source Material | Processed Form | Study type | Model of Infertility | Summary | Reference |
| --- | --- | --- | --- | --- | --- | --- |
| Ovarian engraftment | Healthy Ovarian Tissue | N/A | In vivo | Yes (Chemotherapy) | Batchvarov et al. use a chemotherapy regimen of busulfan and cyclophosphamide to induce infertility. Grafted ovarian tissue was able to rescue host fertility although non-grafted hosts also gave rise to litters 200+ days post treatment. | 70 |
| Biomaterial-facilitated | Sodium Alginate | Alginate gel | In vivo | No | Rios et al. examine the potential of immunisolating and encapsulating ovarian follicles within alginate hydrogels for subcutaneous transplant. Hydrogels were retrieved and mature oocytes were collected then successfully fertilized with 7% of the embryos reaching 4 cell stage. | 17 |
|  | Human Ovary | OECM Scaffold | In vitro/ In vivo | No | Hassanpour et al. develop a decellularization protocol incorporating SLES to prepare an ovarian tissue specific scaffold. | 71 |
|  | Gelatin | 3D Printed Scaffold | In vivo | Yes (Ovariectomy) | Laronda et al. establish a bioprosthetic 3D printed ovary to restore ovarian function in sterilized mice. GFP pups born from exogenous follicle transplant. | 16 |
|  | Synthetic PEG | PEG hydrogel | In vivo | Yes (Ovariectomy) | Kim et al. implement a PEG-VS hydrogel to support the transplantation of immature follicles and restore endocrine function for up to 60 days. | 36 |
|  | Fibrinogen/ Thrombin | Fibrin gel | In vivo | No | Paulini et al. test an artificial ovary using a fibrin gel showing the efficacy of xenografting isolated preantral follicles within the peritoneum. Results showed follicles survival and proliferation up to 7 days post-transplant. | 7 |
|  | Human/ Bovine Ovary | OECM Scaffold | In vivo | Yes (Ovariectomy) | Laronda et al. detail a procedure for the decellularization of both human and bovine ovaries. Primary ovarian cells seeded onto decellularized grafts were transplanted into ovariectomized mice and showed evidence of initiating puberty. | 67 |
|  | Fibrinogen/ Thrombin | Fibrin gel + VEGF | In vitro/ In vivo | Yes (Ovariectomy) | Kniazeva et al. test several biomaterial graft systems in an ovariectomized mouse. Successful litters resulted solely from fibrin gel supplemented with VEGF. | 15 |
|  | Fibrinogen/ Thrombin | Fibrin gel | In vivo | Yes (Ovariectomy) | Smith et al. characterize the use of a fibrin gel for the transplant of ovarian follicles. Early-stage follicles were enzymatically isolated, encapsulated within fibrin gels and transplanted into an ovariectomized mouse. Tissue explants were performed at 0, 3, 9 and 21 days with follicles populations being significantly reduced by day 9. | 72 |
|  | Fibrinogen/ Thrombin | Fibrin gel | In vivo | Yes (Ovariectomy) | Luyckx et al. prepare an artificial ovary from fibrin gel that promotes survival and proliferation of preantral follicles 1 week post-transplant. | 73 |
|  | Fibrinogen/ Thrombin | Fibrin gel + VEGF | In vivo | Yes (Ovariectomy) | Shikanov et al. introduce a combination system consisting of Fibrin gel supplemented with VEGF to stimulate angiogenesis and promote graft survival. Transplanted mouse ovarian tissues supported by the Fibrin/VEGF material improved follicle survival when compared to no biomaterial control. The transplant material was also able to give rise to a litter of pups. | 18 |
| Intraovarian Transplantation | Female germline stem cells (FGSCs) | PBS | In vivo | Yes (Chemotherapy) | Xiong et al. implement a stem cell therapy to reverse the effects observed with exposure to chemotherapy. FGSC microinjection led to healthy pups post-chemotherapy. | 65 |
|  | Primordial Follicles | PBS | In vivo | Yes (Chemotherapy) | Park et al. show significant genetic changes within primordial follicle markers after exposure to chemotherapy. Intraovarian primordial follicle transplantation did not restore fertility. | 64 |

IVM consists of the isolation and culture of immature follicles to obtain meiotically-competent oocytes for IVF. IVM approaches have predominantly shifted from two-dimensional culture toward three-dimensional hydrogel-based follicle encapsulation, which has improved follicle morphology and intercellular signaling^51^. The most commonly used hydrogel for IVM is alginate^9,52–58^; although, there are several other options that have been examined including, fibrin^59^, fibrin-alginate^18,60^, and polyethylene glycol (PEG)^61,62^. Each of these materials provide a unique set of physical and biochemical properties, which allows them to support the growth and maturation of follicles *in vitro*. Successful application of IVM has been shown in mice leading to live-births^57^; however, the pre-clinical translation of this approach for human follicles has been limited^63^. Recently, follicle maturation has been attempted in vivo through the heterotopic subcutaneous transplant of a retrievable hydrogel seeded with immature follicles^17^. Antral follicles developed in the hydrogel and germinal vesicle stage oocytes could be extracted, matured to MII stage and fertilized, leading to the development of two and four cell embryos. However, embryos were not transferred in this study and pregnancies were not established.

As IVM has proven to be a major challenge for human follicles, several groups have pursued the development of an artificial ovary. This concept involves the isolation and sequestering of immature follicles in a bio-supportive scaffold that can be transplanted to recover ovarian function. Similar to IVM, various biomaterials are being examined as options to support the delivery, survival and function of ovarian follicles in vivo. Recently, a fibrin gel supplemented with vascular endothelial growth factor (VEGF) was used to facilitate the transplantation of primordial follicles into the bursa of ovariectomized mice and gave rise to a healthy litter of pups^15^. In another study, a 3D printed gelatin scaffold was used to examine the effects of pore geometry on follicle survival and achieved healthy pups through natural mating post-implantation in sterilized mice^16^. While these methods have shown early success in an ovariectomy model of infertility, it is difficult to gauge the impact that chemotherapy would have on both implant integration and follicle survival. Moreover, these techniques require *in vitro* manipulation of follicles in preparation of the implants in addition to an invasive surgical transplantation procedure, which could have deleterious effects on reproductive outcomes.

Alternatively, intraovarian transplantation has been performed using isolated primordial follicles, without a supportive biomaterial, in a chemotherapy-induced infertility model; however, follicles underwent apoptosis and were unable to rescue fertility^64^. Others have reported establishing female germline stem cells that were transplanted and led to live births in chemotherapy-induced infertile mice^65^. Although these approaches present an intriguing option, they require a significant amount of *ex vivo* cell processing prior to transplant. Additionally, the existence and developmental potential of ovarian stem cells in the adult ovary remains a controversial topic^66^.

To address these concerns and limitations, we have bioengineered an ISO using an OECM hydrogel to support intraovarian follicle delivery. OECM hydrogels were prepared from decellularized porcine ovarian tissues in an attempt to mimic a natural follicle niche. Decellularization has become a common technique to obtain acellular scaffolds with tissue-specific components. Recently, bovine ovaries have been decellularized using detergent-based methods to obtain a tissue-specific scaffold, which was re-seeded with healthy follicles and transplanted to restore cycling in an ovariectomized mouse^67^. Similarly, porcine and bovine decellularized ovaries were further processed into “tissue papers”, demonstrating a versatile approach for short-term *in vitro* culture of nonhuman primate and human ovarian cortical strips^68^. Although acellular ovarian scaffolds have been shown to support follicle function in this form, they are limited by the quantity of follicles that can be seeded and require additional time for *in vitro* culture to allow follicles to attach.

The objective of the present study was to develop a minimally invasive method to deliver immature follicles into the ovary with reduced *in vitro* manipulation and without the requirement of tissue culture. This was accomplished by solubilizing acellular ovarian scaffolds to create an injectable material which was able to reform following injection under physiological conditions. In addition, we proposed an adapted method for efficiently isolating follicles to reduce the total time *ex vivo* prior to transplantation. To mimic ciPOF experienced in the clinic, a single intraperitoneal injection of busulfan and cyclophosphamide was used to significantly reduce the endogenous follicle population in female recipient mice. The OECM hydrogel combined with freshly isolated GFP^+^ ovarian follicles were successfully delivered into the ovarian cortex, forming an ISO. Intraovarian follicle transplant aided by the OECM hydrogel gave rise to multiple, consecutive litters containing at least one pup expressing GFP.

This study demonstrates a potential therapy for restoring fertility after chemotherapy, but we acknowledge that there are several limitations. A major benefit of isolating follicles from the ovarian stroma is the potential to reduce any malignant cells that could be introduced during transplantation. Although we did not validate this with our isolation methods, there have been reports suggesting that follicle isolation can significantly decrease cancer cells prior to implantation^15,69^. In addition, our isolated ovarian cell suspensions were not filtered to contain follicles alone. Therefore, it is possible that stromal cells or other cell types may provide additional benefit for the support of ovarian follicles within the ciPOF ovaries. In addition, donor follicles were isolated from whole allogeneic ovaries rather than autologously sourced ovarian tissue. Furthermore, the ciPOF model used did contain residual endogenous follicles, which could aid exogenous follicle integration and survival within the ovarian niche. However, the effects of chemotherapy on fertility outcomes are both dosage and patient dependent, which our model indicates. The size of mouse ovaries also restricted us from using a larger volume of injected gel during a single treatment making it difficult to deliver a greater quantity of follicles. This limitation is significant, as a higher number of follicles injected would greatly increase the potential for successful fertility outcomes. In addition, the ratio of gel to follicles may also be a decisive factor in long-term follicle survival. For example, a higher follicle concentration could inhibit access to nutrients within the ISO triggering apoptosis or atresia. To this end, a future study would need to be conducted in a large animal, with follicles isolated from cryopreserved, autologous ovarian tissues to adequately evaluate the therapeutic and translational potential of this approach.

OECM hydrogels, paired with intraovarian follicle microinjection, offer a cell delivery platform for fertility preservation. This therapeutic approach employs the innate remodeling capacity of the ECM to establish an ISO to support follicle transplant. The primary advantage of this strategy would provide a minimally invasive surgical intervention akin to transvaginal oocyte retrieval used for IVF. In addition, this approach has the potential to eliminate growth factor supplementation and reduce *in vitro* follicle manipulation. Overall, the restorative reproductive outcomes observed in this study suggest that these alternative methods could address current unmet clinical needs.

## Acknowledgements

We would like to acknowledge funds provided from an anonymous donor and Magee-Womens Research Institute to Dr. Orwig. Also, thanks to the lab animal resource staff of Magee-Womens Research Institute. Special thanks to the Center for Biologic Imaging (CBI) at the University of Pittsburgh for their expertise and use of equipment for obtaining both SEM and multi-photon images.

## Author Contributions

M.J.B., B.N.B and K.E.O. contributed to the study design and prepared the manuscript. M.J.B. developed the protocol for ovarian tissue decellularization, prepared OECM hydrogels, performed characterization of decellularized ovarian tissues (DNA, HYP, sGAG) and hydrogels (rheology and turbidimetric testing), isolated follicles and prepared follicle suspensions for in vivo studies, performed immunofluorescent staining and imaging of transplant recipient ovaries and genotyped litters. M.S. performed follicle microinjection surgeries. K.E.O, M.S. and S.R.S. designed the breeding strategy and performed all mating studies. A.I. and A.L.N. performed protein extractions and ELISAs. M.J.B., Z.X. and S.D. validated antibodies and performed IHC staining and imaging. S.D. performed SEM imaging of ovarian tissues and hydrogels. M.J.B and A.D. analyzed SEM images of OECM hydrogels to assess fiber network characteristics. Z.W.C. performed staining, imaging and follicle quantification of ciPOF ovaries. All authors contributed to the final editing and approval of the manuscript.

## Methods

### Ovarian tissue decellularization

Porcine ovaries from adolescent pigs (< 1 year old) were obtained from the local abattoir (*Thoma Meat Market, Saxonburg, PA*) and immediately stored on ice and frozen at −20°C. Ovaries were thawed, cleared of surrounding connective and adipose tissues, diced into cubes (~0.125 cm^3^) and transferred to a flask containing cold Milli-Q water (MQ). The diced tissues were shaken manually with MQ until residual blood was visibly removed then replaced with fresh MQ and stored overnight at 4°C. The tissues were then rinsed in fresh MQ on an orbital shaker for 30 minutes at 300 rpm. The flask containing tissue was then replaced with a pre-warmed solution of 0.02% trypsin and 0.05% EDTA then agitated on a magnetic stir plate for one hour at 37°C. Ovaries were then rinsed three times with MQ for 15 minutes each at 300 rpm. A 3% solution of Triton X-100 was then added to the flask and shaken for one hour at 300 rpm. A subsequent wash cycle was implemented to remove any residual detergents from the tissues. Each wash cycle consisted of several distilled water rinses with manual shaking (until no bubbles were observed), then alternating washes of MQ and 1X PBS to neutralize and release detergent that was bound to the tissues. After the wash cycle was completed, the fluid was replaced with fresh MQ and stored overnight at 4°C. A 4% sodium deoxycholate solution was added to the flask and agitated for 1 hour at 300 rpm. A subsequent wash cycle was performed then the tissue was replaced with fresh MQ and stored overnight at 4°C. The ovarian tissues were then depyrogenated and disinfected with a 0.1% peracetic acid and 4% ethanol solution for two hours at 300 rpm. This step was followed by three rinses in MQ, 1X PBS, MQ for 15 minutes each at 300 rpm then stored in fresh MQ at 4°C overnight. To ensure adequate removal of detergents and other chemical reagents one final series of washes with MQ, 1X PBS, MQ were performed for 15 minutes each at 300 rpm. The decellularized tissues were then stored at −80°C prior to lyophilization.

### Characterization of decellularized tissues

Decellularized ovarian tissues were characterized using a number of qualitative and quantitative measures to verify the removal of genetic material and maintenance of ovarian specific proteins. Native and decellularized ovarian tissues were formalin-fixed, paraffin-embedded, sectioned and stained using several histological procedures including DAPI (4’,6-diamidino-2-phenylindole), Hematoxylin and Eosin (H&E), and Periodic Acid-Schiff (PAS). Antibodies specific for ECM proteins Collagen I (Abcam, ab34710) and IV (Abcam, ab6586), Fibronectin (Abcam, ab23751), and Laminin (Abcam, ab11575) were evaluated using DAB (3,3’-Diaminobenzidine) immunohistochemistry (IHC) staining to show conservation after decellularization. IHC tissue sections were counterstained using Hematoxylin QS (Vector Labs, Cat No. H-3404) to show cell nuclei in contrast with resident ECM proteins. DNA removal was quantified using a PicoGreen dsDNA assay kit (Invitrogen, Cat No. P11496). A 2.5% agarose gel was used to detect DNA fragments at a resolution between 25 and 1000 bp. Hydroxyproline (HYP) and sulfated glycosaminoglycans (sGAG) assays were performed to detect collagen and sGAG content. Native and decellularized ovarian tissues were homogenized in a High Salt Buffer (pH 7.5, 50 mM Tris base, 150 mM NaCl, 5 M CaCl_2_, 1% Triton-X-100, 1% Halt protease inhibitor cocktail, Pierce Biotechnology, Rockford, IL). Protein concentrations of the extracted tissues were determined using the BCA Protein Assay Kit (Pierce Biotechnology, Rockford, IL) and 50 ug total protein was used per sample for all assays. Ovarian specific growth factors including insulin growth factor (IGF-1) (R&D Systems, Minneapolis, MN), 17β-estradiol (R&D Systems, Minneapolis, MN), progesterone (Abcam, Cambridge, MA), anti-Müllerian hormone (AMH) (R&D Systems, Minneapolis, MN), and vascular endothelial growth factor (VEGF) (Abcam, Cambridge, MA), were quantified using enzyme-linked immunosorbent assays (ELISA).

### Ovarian ECM digestion and hydrogel formation

Lyophilized ovarian ECM powder was solubilized via enzymatic digestion. A stock ECM digest concentration of 10 mg/mL was prepared by adding 200 mg of ECM powder to a 20 mL solution of pepsin (Sigma P7012) at a concentration of 1 mg/mL (≥2,500 units/mg) dissolved in 0.01 N hydrochloric acid (HCl). Digestion was facilitated with an overhead mixer between 700-2000 RPM for less than 48 hours. Hydrogels were formed after neutralizing and buffering the solubilized ovarian ECM to physiological conditions. Two hydrogel concentrations (4 and 8 mg/mL) were prepared for testing and experimentation. A pre-gel solution was made on ice using the following components: (i) 10 mg/mL OECM digest stock (volume determined by desired final concentration) (ii) 0.1 N NaOH (1/10^th^ the volume of the digest), (iii) 10X PBS (1/9^th^ the volume of the digest), and (iv) 1X PBS or L-15 medium (brought up to final volume). The solution was pulsed 3 times on a vortexer to mix then stored at 4°C until further use.

### Hydrogel characterization

Ovarian hydrogel ultrastructure was assessed using scanning electron microscopy (SEM). Hydrogels were fixed using 2.5% glutaraldehyde, washed with 1X PBS and post-fixed with osmium tetroxide (OsO_4_). Samples were washed again in 1X PBS to dilute the OsO4 then they were slowly exsiccated through a series of increasing ethanol concentrations. Complete dehydration was achieved using a critical point dryer. Dried hydrogels were sputter-coated with gold/palladium particles and imaged at 8,000X magnification. A proprietary Matlab code was used to analyze SEM images to determine various hydrogel fiber network characteristics. Ovarian hydrogel bulk viscoelastic properties were determined using a dynamic parallel plate (40 mm) rheometer (AR-2000 TA instruments). A time sweep (5% strain, 1 rad/s) was used to demonstrate the effect of ECM concentration on both the storage (G’) and loss (G”) modulus. Turbidimetric gelation kinetics were performed on the ovarian hydrogels as previously described^74,75^.

### Chemotherapy-induced POF model

Busulfan (Sigma) and cyclophosphamide (MP BioMedicals LLC) were combined to induce POF in 6-week old female mice (NCR nu/nu). Recipient mice were given a single intraperitoneal injection (IP) and allowed to recover up to 3 weeks prior to treatment. To initially identify the most appropriate chemotherapy regimen, four doses of busulfan/cyclophosphamide (mg/kg) were tested: (1) 12-100 (2) 12-200 (3) 24-100 (4) 24-200. Ovaries were excised, fixed in 4% paraformaldehyde (PFA), paraffin embedded and serial sectioned. Tissue sections were stained using Weigert’s Hematoxylin Picric acid Methyl Blue then imaged under an upright bright field microscope.

Every 10 sections were examined for total follicle number and classified by stage and quantified. The following criteria was used to count the follicles: (1) each follicle contains a visible oocyte (2) Primordial follicles have a single layer of squamous granulosa cells (3) Primary follicles have single layer of cuboidal granulosa cells (4) Secondary follicles contain two to four layers of granulosa cells without the development of an antrum (5) Antral follicles have greater than four layers of granulosa cells as well as definitive antrum. The total number of follicles were quantified then the sum was multiplied by 10 to provide an estimate of the entire follicle population of each ovary.

### Enzymatic follicle isolation

Ovarian donor follicles were prepared using a physical and enzymatic isolation procedure adapted from Kim et al.^36^. First, ovaries were excised from 6-14 day old female (DBA GFP/nu) mice and placed into pre-warmed L-15 (Leibovitz’s) medium. The ovaries were freed from the bursa using a pair of forceps and an insulin needle. Ovaries were then minced using insulin needles into small fragments to aid in digestion. The ovarian fragments were then added to 1.5 mL microcentrifuge tube containing 500 μL of pre-warmed L-15 medium and 50 μL of Liberase TM (13 Wünsch units/mL). The tubes were then placed on an orbital shaker and agitated at 200 rpm for 5 minutes at 37°C. After incubation, the mixture was then pipetted gently for one minute to help free the ovarian follicles from connective tissue. This process was repeated once more until the ovaries had been completely dissociated. After digestion, 10% fetal bovine serum was added to the mixture to halt enzyme activity. The tubes were placed in an upright position for 15 minutes at 37°C to allow follicles to sediment. After 15 minutes, 200 μL of the mixture was carefully pipetted off the top to remove singular ovarian cells. The samples were centrifuged at 100*g* for 5 minutes to loosely concentrate the follicles. A syringe needle was used to gently remove the medium from the tube without disturbing the follicle pellet. Finally, the pellet was resuspended in a chilled 4 mg/mL OECM pre-gel and kept on ice in preparation for follicle microinjection.

### In vivo follicle microinjection

Eight-week old ciPOF female mice (NCR nu/nu) mice were anesthetized and placed on the operating table with their back exposed. A single midline dorsal incision (0.5 cm) was made using small scissors. Subcutaneous connective tissue was then freed from the underlying muscle on each side using blunt forceps. Once the ovary was located under the thin muscle layer, a small incision (<1 cm) was made to gain entry to the peritoneal cavity. The edge of the incision was held with tooth forceps, while the ovarian fat pad was removed to expose the ovary and surrounding bursa. A small volume (<10 μL) of chilled follicle-OECM pre-gel was transferred to a glass needle (filament Cat#: FB245B and borosilicate glass micro pipette Cat#: B100-75-10) then secured to a pressurized microinjection system. Eppendorf microinjection system (TransferMan NK2 and FemtoJet) was used for follicle delivery and surgical manipulation observed under a Nikon SMZ stereomicroscope. The loaded needle was positioned perpendicular to the ovary and guided into the ovarian cortex, where the follicle-OECM mixture was slowly injected at a constant pressure ranging from 50-250 hPa. For each injection, a minimum of one ovary worth of follicles was transplanted. The same surgical and injection procedures were performed contralaterally. The follicle-injected ovaries were placed back into the abdominal cavity, the muscle layers were sutured, and the skin incision was stapled.

### Mating Study

Two weeks after injection both the follicle recipient and non-injected control ciPOF nude female mice were bred to male nude mice (NCR nu/nu, Taconic). The breeding was conducted for three cycles, which concluded at 106 days on average. Pups born were fostered within 1 day with NCR nu/+ (Taconic) females due to the lack of developed mammary glands in the nude mouse strain used for recipients, then they were weaned at 3-4 weeks old. Pups (DBA-GFP/nu-, Orwig Lab) inherited from the follicle injected recipients were selected based on physical traits consisting of fur, dark eyes or GFP expression. Genotyping was performed using mouse tail DNA and standard PCR with the following primers: GFP forward primer sequence (GAA CGG CAT CAA GGT GAA CT); GFP reverse primer sequence (TGC TCA GGT AGT GGT TGT CG); β-actin forward primer sequence (CGG TTC CGA TGC CCT GAG GCT CTT); β-actin reverse primer sequence (CGT CAC ACT TCA TGA TGG AAT TGA) (primers prepared by Integrated DNA Technologies, inc.). PCR products were run on a 2.5% agarose gel and imaged under UV light. The resulting pups were grown to 8 weeks and bred for fertility status. Second generation breeding pairs consisted of a GFP/nu-experimental female and DBA/2 control male (Jackson), a GFP/nu-experimental female and GFP/nu-experimental male, and a DBA/2 control female and GFP/nu-experimental male. Breeding pairs were separated after two weeks.

### Statistical analysis

All data were expressed as mean ± s.e.m and plotted using GraphPad Prism 7.02. For the analysis of normally (parametrically) distributed data, the individual means were compared using an unpaired, two-tailed, t-test. For the analysis of non-parametrically distributed data, the mean ranks were compared using an unpaired, one-way ANOVA (Kruskal-Wallis) with adjusted P-values calculated based upon Dunn’s multiple comparisons test. Exact P-values resulting from the statistical analyses are presented within each figure.

