## Supplemental Figures for "Bioengineering an *in situ* ovary (ISO) for fertility preservation"

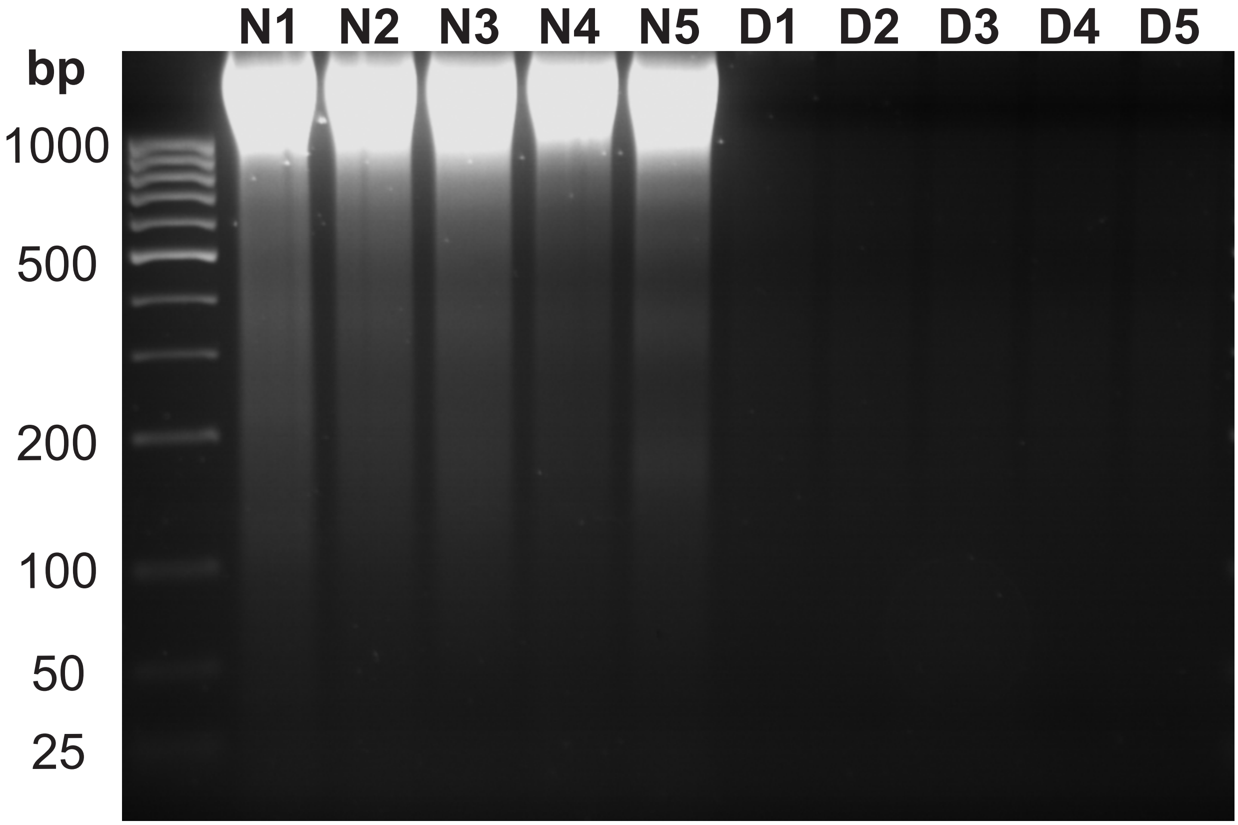

**Supplemental Fig. 1| Assessment of DNA Removal with Agarose Gel Electrophoresis.** A 2.5% agarose gel was used to characterize the presence of DNA within ovarian tissues post-decellularization. No visible bands were observed within the decellularized ovarian tissues (D1-D5). Native ovarian tissues (N1-N5) were used as a positive control. Each lane represents an individual sample of DNA.

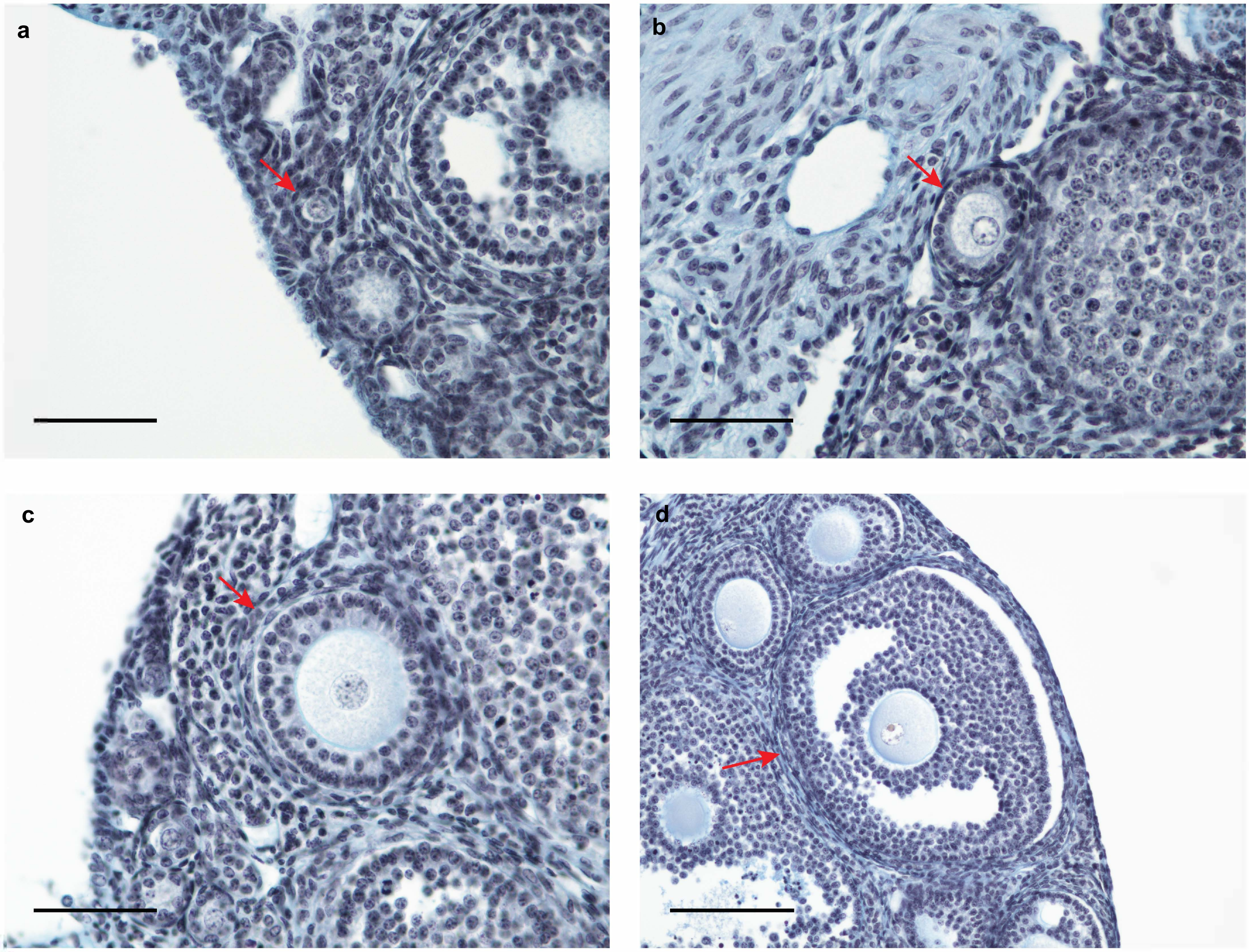

**Supplementary Fig. 2| Follicle stage characterization.** Follicles were quantified and developmental stage was determined by their morphology **a**, Primordial follicles were recognized by a central oocyte surrounded by a single layer of squamous granulosa cells. Scale, 50 µm. **b**, Primary follicles were counted if they contained a single oocyte with a layer of cuboidal granulosa cells. Scale, 50 µm. **c**, Secondary follicles contained an oocyte with 2-4 layers of cuboidal granulosa cells. Scale, 50 µm. **d**, Antral follicles were distinguished by an oocyte with several layers of cuboidal granulosa cells containing pockets of antral fluid. Scale, 100 µm. Red arrows indicate counted follicles.

**a**

| | | Female | | | |
|---|---|---|---|---|---|
| | | **Endogenous Nude Mouse Follicles** | | **Transplanted GFP Mouse Follicles** | |
| | | Nu- | Nu- | DBA GFP+ | DBA GFP- |
| Male | Nu- | NCR nu-/nu-: nude mice with red eyes | NCR nu-/nu-: nude mice with red eyes | GFP+/nu-: furry, dark eyes, glows green under UV | GFP-/nu-: furry, dark eyes, does not glow green |
| | Nu- | NCR nu-/nu-: nude mice with red eyes | NCR nu-/nu-: nude mice with red eyes | GFP+/nu-: furry, dark eyes, glows green under UV | GFP-/nu-: furry, dark eyes, does not glow green |

**b**

| | | Cycle #1 | | | Cycle #2 | | | Cycle #3 | | | | |
|---|---|---|---|---|---|---|---|---|---|---|---|---|
| Chemotherapy Dose | Animal ID | Live | Dead | GFP | Live | Dead | GFP | Live | Dead | GFP | Total Pups | Total GFP |
| 12-100 | 6194 | 4 | 0 | 0 | 0 | 0 | 0 | 0 | 0 | 0 | 4 | 0 |
| | 6195 | 2 | 4 | 0 | 3 | 0 | 0 | 5 | 0 | 1 | 14 | 1 |
| | 6196 | 2 | 0 | 0 | 1 | 4 | 0 | 0 | 0 | 0 | 7 | 0 |
| | 6197 | 0 | 0 | 0 | 3 | 0 | 0 | 6 | 0 | 0 | 9 | 0 |
| | 6198 | 0 | 0 | 0 | 0 | 0 | 0 | 0 | 0 | 0 | 0 | 0 |
| 24-100 | 6237 | 0 | 0 | 0 | 0 | 0 | 0 | 0 | 0 | 0 | 0 | 0 |
| | 6238 | 3 | 0 | 1 | 1 | 0 | 1 | 0 | 0 | 0 | 4 | 2 |
| | 6240 | 11 | 0 | 0 | 5 | 0 | 0 | 0 | 0 | 0 | 16 | 0 |
| | 6241 | 0 | 0 | 0 | 0 | 0 | 0 | 0 | 0 | 0 | 0 | 0 |

**c**

| | GFP+/Nu+ | GFP+/Nu- | GFP-/Nu+ | GFP-/Nu- |
|---|---|---|---|---|
| GFP-/Nu+ | GFP+/- Nu+/+ | GFP+/- Nu+/- | GFP-/- Nu+/+ | GFP-/- Nu+/- |
| GFP-/Nu+ | GFP+/- Nu+/+ | GFP+/- Nu+/- | GFP-/- Nu+/+ | GFP-/- Nu+/- |
| GFP-/Nu+ | GFP+/- Nu+/+ | GFP+/- Nu+/- | GFP-/- Nu+/+ | GFP-/- Nu+/- |
| GFP-/Nu+ | GFP+/- Nu+/+ | GFP+/- Nu+/- | GFP-/- Nu+/+ | GFP-/- Nu+/- |

**d**

| | GFP+/Nu+ | GFP+/Nu- | GFP-/Nu+ | GFP-/Nu- |
|---|---|---|---|---|
| GFP+/Nu+ | GFP+/+ Nu+/+ | GFP+/+ Nu+/- | GFP+/- Nu+/+ | GFP+/- Nu+/- |
| GFP+/Nu- | GFP+/+ Nu+/- | GFP+/+ Nu-/- | GFP+/- Nu+/- | GFP+/- Nu-/- |
| GFP-/Nu+ | GFP+/- Nu+/+ | GFP+/- Nu+/- | GFP-/- Nu+/+ | GFP-/- Nu+/- |
| GFP-/Nu- | GFP+/- Nu+/- | GFP+/- Nu-/- | GFP-/- Nu+/- | GFP-/- Nu-/- |

**e**

| Animal ID | Breeding Partner ID | Cycle | Live | Dead | GFP |
|---|---|---|---|---|---|
| **GFP/nu-001 (F)** | DBA - 1683 (M) | 1 | 10 | 0 | 5 |
| | **GFP/nu-003 (M)** | 2 | 8 | 1 | 6 |
| **GFP/nu-002 (F)** | **GFP/nu-003 (M)** | 1 | 10 | 0 | 9 |
| | DBA - 1684 (M) | 2 | 0 | 0 | 0 |
| DBA/2-1644 (F) | B6D2 - 853 (M) | 1 | 11 | 0 | 0 |
| | **GFP/nu-003 (M)** | 2 | 8 | 0 | 6 |
| | **GFP/nu-003 (M)** | 3 | 7 | 0 | 3 |
| **Total Second Generation Pups** | | | 43 | 1 | 29 |

**Supplementary Fig. 3| Breeding Strategy and Results. a**, The table represents the potential breeding outcomes expected after intraovarian follicle microinjection. Genetic backgrounds may be derived from both endogenous and transplanted follicles. Pups born from the transplanted follicles were distinguished by GFP expression and/or fur with dark eyes. **b**, Three cycles of breeding occurred with the tranplanted ciPOF nude (nu-/nu-) female mice mated to nude (nu-/nu-) male mice. Three GFP pups were born as a result of the follicle transplation. Non-injected control ciPOF mice are highlighted in red. Pups generated from the follicle transplant were bred to determine reproductive health. Potential breeding outcomes shown for **c**, GFP/nu pups bred with DBA wild-type (GFP-/nu+) mice and **d**, GFP/nu pups bred with GFP/nu pups. Cells outlined in red indicate nude offspring (nu-/nu-). **e**, Multiple breeding cycles with the follicle-transplant derived GFP mice resulted in a total of 43 second generation pups.

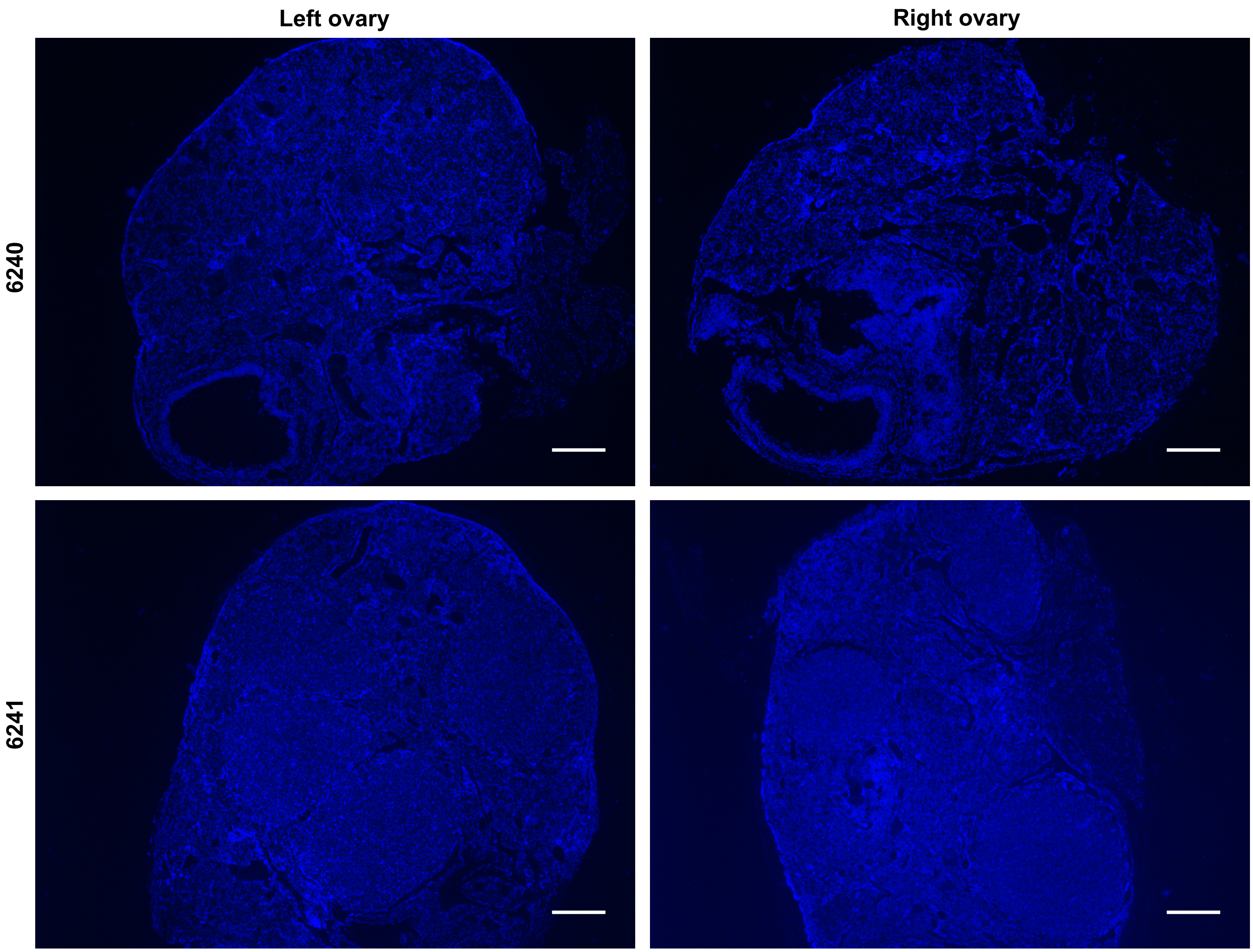

**Supplementary Fig. 4| Non-injected ciPOF control ovaries.** Merged DAPI and FITC immunofluorescence images of control ovaries (Animal ID: 6240 and 6241) Scale, 200 µm. Non-injected control tissues showed endogenous follicle growth, but appeared to have a reduced population of immature follicles in comparison to ovaries that received follicle transplant. As expected, no cells within the control tissues expressed GFP.
